# Integrating Narrow-Window DIA with AI-Powered Search Enables Deep and Reliable Functional Proteomics

**DOI:** 10.64898/2026.09.04.749476

**Authors:** Yun Xiong, Dejan Stepec, Huimin Zhang, Lin Tan, Bo Wei, Maximilien Burq, John N. Weinstein, Peter Cimermancic, Philip L. Lorenzi

**Affiliations:** Department of Bioinformatics and Computational Biology, The University of Texas MD Anderson Cancer Center (MDACC), Houston, TX 77030, USA; Proteomics Core Facility, Department of Bioinformatics and Computational Biology, The University of Texas MD Anderson Cancer Center (MDACC), Houston, TX 77030, USA; Metabolomics Core Facility, Department of Bioinformatics and Computational Biology, The University of Texas MD Anderson Cancer Center (MDACC), Houston, TX 77030, USA; Department of Experimental Radiation Oncology, The University of Texas MD Anderson Cancer Center, Houston, TX 77030, USA; Tesorai Inc, USA; Department of Systems Biology, The University of Texas MD Anderson Cancer Center (MDACC), Houston, TX 77030, USA; Microbiome Program, Department of Hematology and Hematopoietic Cell Transplantation, City of Hope National Medical Center, Duarte, CA 91010, USA

## Abstract

Narrow-window data-independent acquisition (nDIA) is emerging as a powerful technique for bottom-up proteomics. Here, we systematically benchmarked nDIA, wide-window DIA (wDIA), narrow-window data-dependent acquisition (nDDA), and wide-window DDA (wDDA) for rapid, single-shot proteomic analysis. For data processing, we introduced Tesorai Search, a new search engine leveraging a large pre-trained model and compared it with DIA-NN and FragPipe across both DIA and DDA datasets. Among 12 acquisition-analysis pipelines evaluated, nDIA combined with DIA-NN and Tesorai Search delivered the highest proteome coverage, identifying 10,255 and 10,766 protein groups from benchmark samples, respectively. Both search engines maintained rigorous false-discovery rate (FDR) control. While nDIA generally outperformed nDDA in sensitivity, FragPipe-DDA+ approach proved to be the most sensitive within the nDDA pipelines. However, entrapment analyses indicate that this sensitivity comes at the cost of less robust FDR control compared to Tesorai Search. As a proof of concept, we applied nDIA-MS to 17 cancer cell lines harboring DNA damage response (DDR) gene knockouts, successfully detecting significant downregulation of all targeted proteins and uncovering 81 DDR-related proteins modulated in at least one cell line. These results underscore nDIA-MS, together with DIA-NN and Tesorai Search, as a robust and scalable platform for high-throughput functional proteomic screening.

## Introduction

Mass spectrometry (MS)-based proteomics has become a cornerstone technology for characterizing proteomes at scale, enabling in-depth profiling of protein abundance and dynamics across a wide range of biological systems and clinical contexts(*1, 2*). In bottom-up proteomics, two principal data acquisition strategies are commonly employed: data-dependent acquisition (DDA) or data-independent acquisition (DIA)(*3–5*). These methods differ fundamentally in how precursor ions are selected and fragmented. In DDA, precursor ions are selected in real-time based on MS1 survey scans, prioritizing the most abundant ions for MS2 fragmentation. In contrast, DIA uses a predefined series of mass isolation windows to fragment all ions within each window simultaneously, removing the need for real-time precursor selection. Whereas DDA offers high precursor selectivity, DIA usually generates highly multiplexed MS2 spectra and complex chromatograms. To mitigate interference from co-eluting precursors, DIA workflows typically segment the m/z range into multiple windows. Among those workflows, wide-window DIA with isolation windows ranging from 10 to 100 Th is commonly adopted due to its ease of implementation and broad instrument compatibility(*3*). However, wide isolation windows can compromise precursor specificity, resulting in ambiguous precursor-fragment relationships.

A promising solution involves using narrow isolation windows (e.g., 2 Th), which improves precursor selectivity but traditionally requires multiple injections or compromised cycle time due to limited instrument scan speed(*3*). As a result, narrow-window DIA (nDIA) has been difficult to implement across a full m/z range in a single injection. The Orbitrap Astral is a next-generation MS platform that enables parallel MS1 and MS2 acquisition and combines high-resolution MS1 analysis at 240,000 mass resolution (at m/z 200) in the Orbitrap with ~200 Hz MS2 scan speed at 80,000 mass resolution in the asymmetric track lossless (Astral) analyzer(*6–8*). The timsTOF family of instruments are also next-generation MS platforms that combine trapped ion mobility spectroscopy (TIMS) resolution of 200 in the TIMS cell with ~300 Hz MS2 scans at up to 60,000 mass resolution in the time of flight (TOF) analyzer. These configurations make single-injection nDIA feasible, unlocking new opportunities for high-throughput, high-specificity proteomics(*3, 9–13*).

In this study, we present a systematic benchmarking of nDIA on the Orbitrap Astral, comparing its performance with conventional wide-window DIA, narrow-window DDA, and wide-window data-dependent acquisition (WWA, or wDDA) strategy(*14*). For data processing, we compared the performance of Tesorai Search(*15*), a new search engine based on a large pre-trained model, and two commonly used software suites, DIA-NN and FragPipe, across both DIA and DDA datasets(*16–18*). To demonstrate its performance, we applied nDIA-MS to protein extracts from 17 cancer cell lines with single knockout of specific DNA damage response (DDR) genes(*19–22*). Our findings support the reliability and utility of nDIA-MS for high-throughput functional proteomic screening and demonstrate its value in large-scale cancer biology research.

## Results

### MS proteomics workflows for peptide and protein identification

Leveraging the capabilities of the Orbitrap Astral mass spectrometry (MS) system, nDIA has been recognized to significantly enhance the capability of single-shot, high-throughput proteomics for complex proteome profiling(*9, 13*). We systematically compared the performance of four data acquisition technologies: nDIA, conventional wide-window DIA (DIA or wDIA)(*23*), narrow-window DDA (DDA, or nDDA), and wide-window acquisition (WWA, or wDDA)(*14, 24*), which is a modified DDA method that uses wide mass isolation windows (Figure 1A).

**Figure 1.**
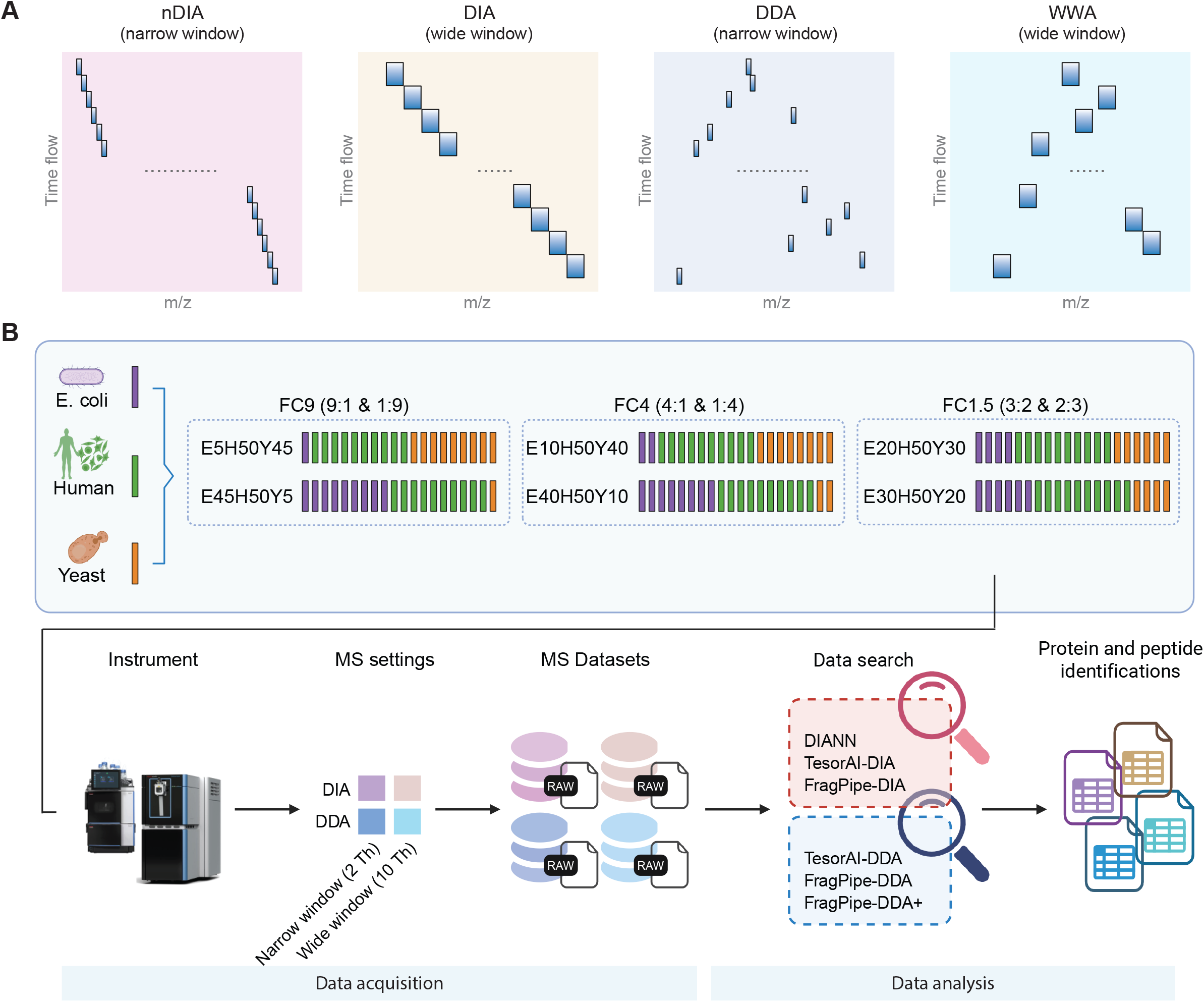
Overview of the benchmark experiments. **(A)** Schematic of the four types of MS data acquisition approaches, including narrow window DIA (nDIA), wide window DIA, classical narrow window DDA, and wide window acquisition (WWA). **(B)** The benchmark sample sets and benchmark experiments workflow. Peptides from E. coli, human, and yeast cells were mixed at different ratios. The mixed three species samples were subjected to Orbitrap Astral MS system equipped with Vanquish Neo HPLC. For each sample, all the four aforementioned data acquisition approaches were used to collect the MS datasets. Raw MS datasets were processed by Tesorai Search, DIA-NN, and FragPipe. Peptide and protein identification and quantitative information were used for downstream analysis.

For data processing, we introduce Tesorai Search(*15*), a search engine based on a large pre-trained model that supports both DDA and DIA workflows, and compare it with two widely used software tools: DIA-NN(*16*), primarily used for DIA analysis, and FragPipe(*25*), which also supports both DDA and DIA workflows (Figure 1B). Notably, FragPipe offers DDA, DIA, and DDA+ workflows(*18*), the latter of which is designed for analyzing wide-window DDA datasets(*24, 25*). Combining the four acquisition strategies and three software tools resulted in 12 distinct acquisition-analysis combinations.

To prepare benchmarking samples, we generated a mixed-species peptide standard consisting of *Escherichia coli*, human, and yeast peptides in six different ratios: 5:50:45 (sample E5H50Y45), 10:50:40 (sample E10H50Y40), 20:50:30 (sample E20H50Y30), 30:50:20 (sample E30H50Y20), 40:50:10 (sample E40H50Y10), and 45:50:5 (sample E45H50Y5) (Figure 1B). Those samples were analyzed on the Orbitrap Astral MS using the four acquisition strategies described above (Figure 1A).

### nDIA with DIA-NN and Tesorai Search yields the deepest proteome coverage

We first performed a comparative statistical analysis of protein and peptide identifications (Figure 2A-D, Supplementary Table 1-6). Among all methods evaluated, nDIA combined with DIA-NN and Tesorai Search yielded the deepest proteome coverage, with an average of 10,255 and 10,766 proteins identified from the benchmark samples, respectively. nDIA/DIA-NN result identified an average of 97,976 peptides, which outperformed Tesorai Search and FragPipe at the peptide level (Figure 2A, C). nDIA/Tesorai Search achieved the second-greatest peptide identification, followed by nDIA/FragPipe. DIA methods consistently outperformed DDA methods in terms of identification depth, regardless of the isolation window size (Figure 2A-D). Moreover, data acquired with a narrow isolation window (2 Th) produced more identifications than data acquired with a wide isolation window (10 Th).

**Figure 2.**
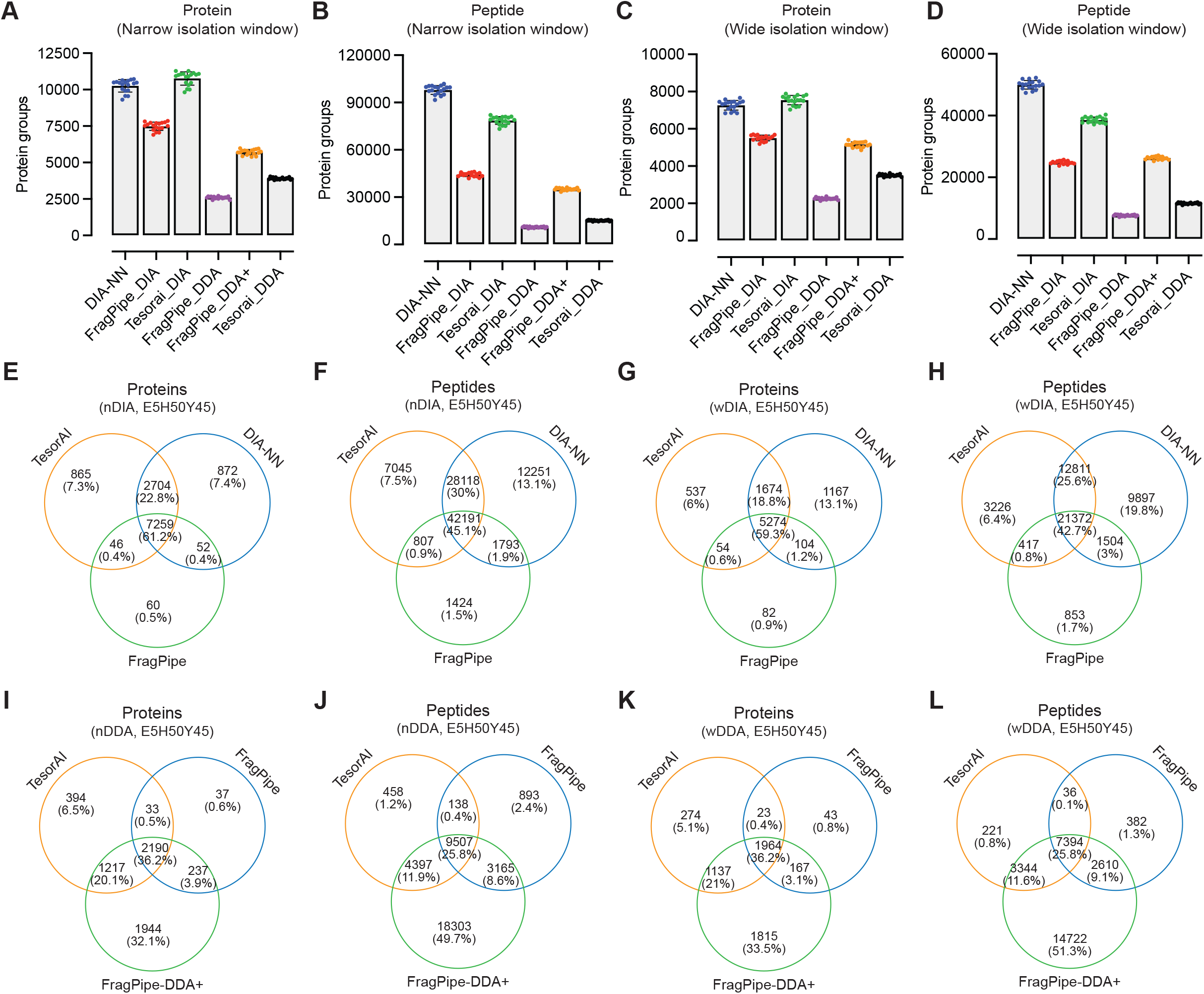
Comparative analysis of the peptides and proteins identification by different software suites from DIA and DDA datasets. **(A-D)** Statistic summary of the number of peptides **(A, B)** and proteins **(C, D)** identified by different strategies. **(E-H)** Overlap analysis of peptides and proteins identified by Tesorai Search, DIA-NN, and FragPipe from DIA datasets generated with narrow **(E, F)** or wide **(G, H)** isolation windows. **(I-L)** Overlap analysis of peptides and proteins identified by Tesorai Search, FragPipe-DDA, and FragPipe-DDA+ from DDA datasets generated with narrow **(I, J)** or wide **(K, L)** isolation windows.

Whereas DIA-NN and Tesorai Search yielded more protein identifications than FragPipe for DIA datasets, with DDA datasets the DDA/FragPipe-DDA+ workflow exhibited roughly twice the protein identification depth of DDA/FragPipe and DDA/Tesorai Search^18^. These results suggest that FragPipe-DDA+ is a promising choice in protein and peptide identification for DDA datasets.

To evaluate overlapping and unique identifications across workflows, we analyzed the overlap of output acquired for the benchmark sample E5H50Y45. With nDIA datasets, 45% and 61% of annotated peptides and proteins, respectively, were commonly identified by Tesorai Search, DIA-NN, and FragPipe (Figure 2E, F). The overlap between Tesorai Search and DIA-NN was 75% for peptides and 84% for proteins. FragPipe and DIA-NN exhibited only 47% overlap for peptides and 61% for proteins. A similar trend was observed in wide window DIA (wDIA) datasets (Figure 2G, H). These results indicate high concordance between Tesorai Search and DIA-NN with DIA data processing.

For DDA datasets generated using a narrow isolation window (nDDA), 26% and 36% of annotated peptides and proteins were commonly identified across Tesorai Search, FragPipe, and FragPipe-DDA+ (Figure 2I, J). The FragPipe-DDA+ approach uniquely identified 50% of peptides and 32% of proteins. Tesorai Search shared 38% of peptides and 56% of proteins with FragPipe-DDA+, representing a higher overlap than observed between FragPipe-DDA+ and standard FragPipe. Wide window DDA (wDIA) datasets yielded a similar pattern (Figure 2K, L). Collectively, these findings demonstrate the robustness of Tesorai Search across both acquisition strategies and underscore a considerable advantage of FragPipe-DDA+ with DDA datasets.

### DDA+ Achieves Significant Overlap with DIA

With both Tesorai Search and FragPipe capable of processing DDA and DIA data, direct comparison of DDA vs. DIA acquisition strategies become feasible. DIA clearly outperforms DDA in peptide and protein identification. According to Tesorai Search result, 82% and 72% of all annotated proteins are specifically identified by DIA with narrow or wide window, respectively (Figure 3A-D). Similar trends were also found at peptide level.

**Figure 3.**
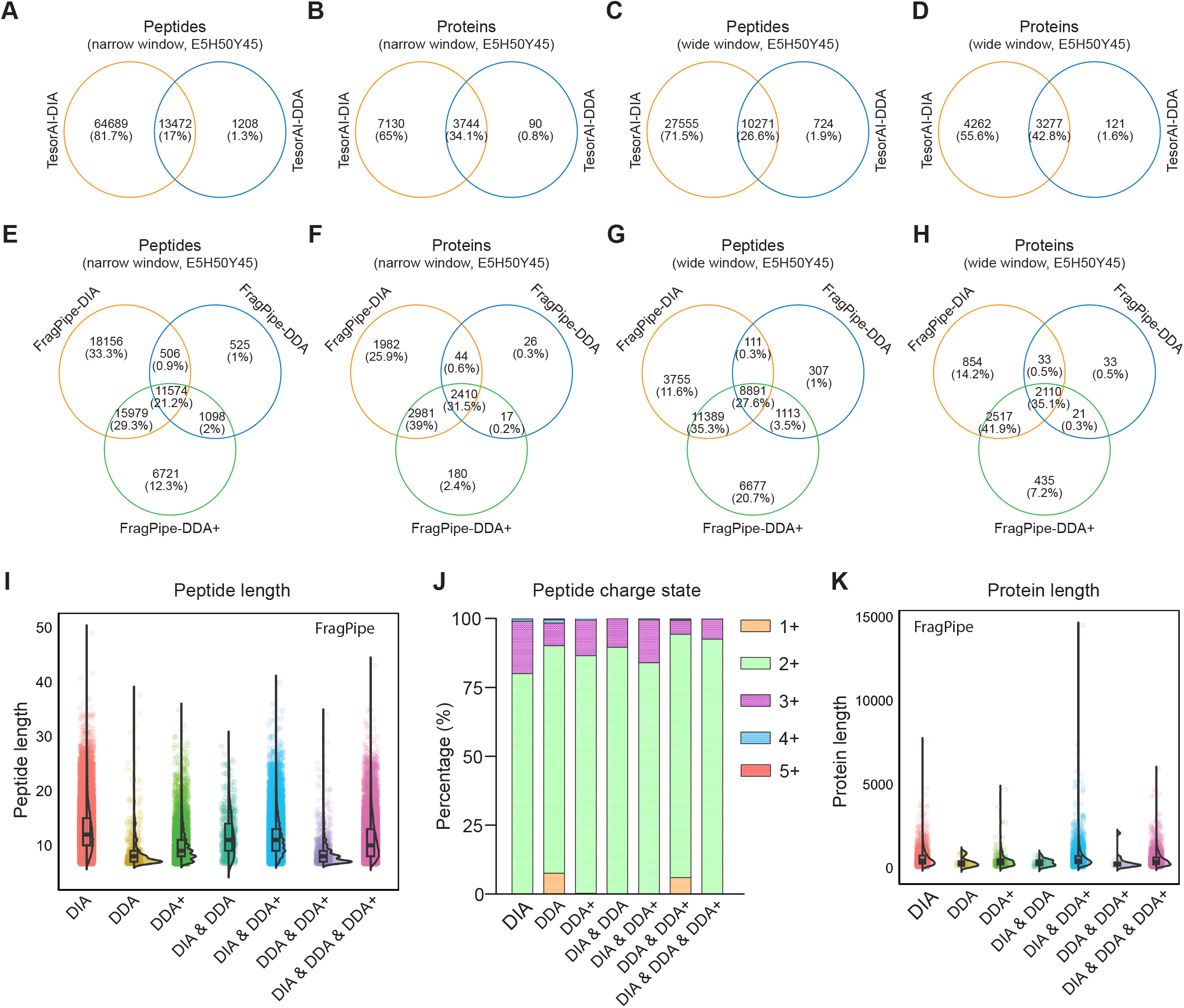
Direct comparison of the peptides and proteins identification by DIA and DDA. **(A-D)** Overlap analysis of peptides and proteins identified in Tesorai Search output by DIA and DDA with narrow **(A, B)** or wide **(C, D)** isolation window. **(E-H)** Overlap analysis of peptides and proteins identified in FragPipe output by DIA and DDA with narrow **(E, F)** or wide **(G, H)** isolation window. **(I)** Distribution of peptide length across DIA, DDA, and DDA+ workflows. **(J)** Distribution of peptide charge states across DIA, DDA, and DDA+ workflows. **(K)** Distribution of protein length across DIA, DDA, and DDA+ workflows.

FragPipe results were consistent with those of Tesorai Search, indicating a clear advantage of DIA acquisition over DDA. Within DDA datasets, though, FragPipe-DDA+ demonstrated a significant improvement over standard DDA. Among the union of all datasets, nDDA/DDA+ uniquely identified 12% of all annotated peptides and 2.4% of all annotated proteins, compared to 1.0% and 0.3% by nDIA/DDA, respectively (Figure 3E, F). This advantage was even more pronounced with wide windows, where DDA/DDA+ uniquely identified 21% of peptides, surpassing all DIA (~12%) and DDA (~1%) workflows (Figure 3G). The enhanced performance of DDA+ under wide window conditions may reflect its ability to better leverage increased spectral complexity, which can hinder the effectiveness of DIA. Under narrow window conditions, 51% of peptides and 71% of proteins were shared between nDIA/FragPipe-DIA and nDDA/FragPipe-DDA+ results (Figure 3E, F). These values increased to 63% and 77%, respectively, with wide window acquisitions (Figure 3G, H). Together, these results suggest the complementary strengths of DIA and DDA+ for data acquisition and analysis and highlight their robustness and reliability for comprehensive proteomic analysis.

### DIA and DDA+ preferentially identify peptides with longer sequences and higher charge states

To elucidate the mechanisms underlying the advantages of DIA and DDA+, we examined the physicochemical properties of peptides and proteins identified uniquely or commonly by different analytical approaches (Figure 3I, J). As illustrated, a greater proportion of longer peptides with higher charge states were identified in the DIA and DDA+ datasets. In contrast, we did not observe an apparent difference in protein length (Figure 3K). We hypothesize that longer peptides with higher charge states may be more prone to co-elution with regular peptides, resulting in spectra of higher complexity. Such spectra, which are typically excluded in standard DDA analysis, are more readily captured and identified by DIA and DDA+.

### Impact of Isolation Window on Peptide and Protein Identification

Although wide window acquisition has been reported to improve protein identification rates in DDA workflows using traditional mass spectrometry instruments(*14, 24*), our DDA datasets produced by Orbitrap Astral did not benefit from the use of wide isolation windows; no significant difference in protein identification was observed between narrow and wide isolation windows with DDA workflows (Figure 2A-D). For DIA workflows, we observed a notable decrease in the number of proteins identified using wide isolation windows compared to narrow isolation windows.

To further investigate the effects of isolation window size, we analyzed protein digestion samples from human HeLa cells across a range of isolation windows from 1 to 20 Th (Figure 4, Supplementary Table 7-12). Consistent with previous reports(*13*), optimal performance for DIA was observed with a 2 Th isolation window. For DDA, the optimal isolation window was 5 Th, particularly when DDA+ mode was employed. With an isolation window greater than 10 Th, the high complexity of mass spectra appears to have suppressed the performance of both DIA and DDA approaches.

**Figure 4.**
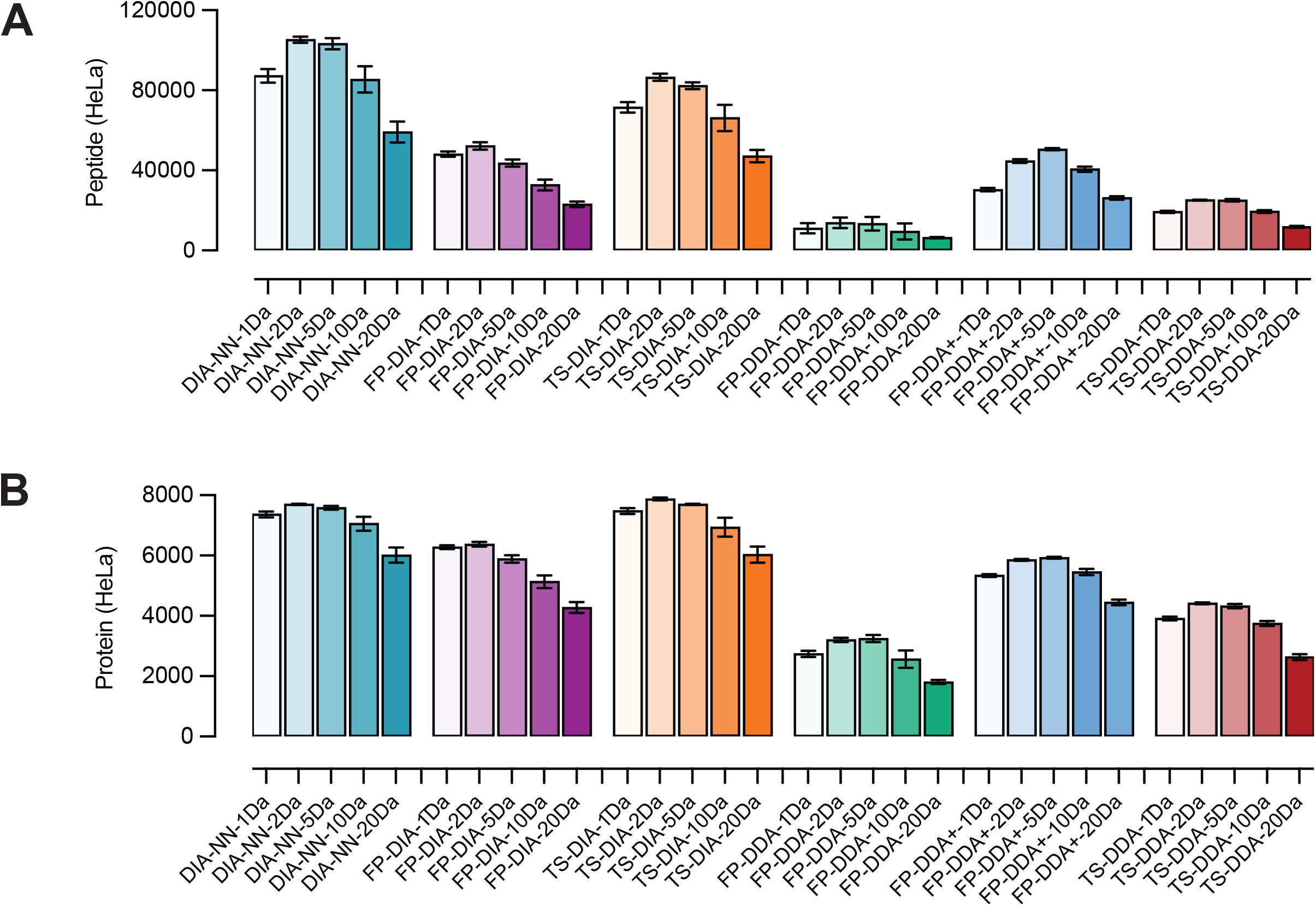
Effect of isolation window range on peptide and protein identification. HeLa digest samples were subjected to MS analysis using DIA or DDA approaches with a variety of isolation windows ranging from 1 Th to 20 Th. The generated DIA datasets were processed by DIA-NN, Tesorai Search, and FragPipe. DDA datasets were processed using Tesorai Search, FragPipe-DDA, and FragPipe-DDA+ methods.

### Entrapment Analysis for Assessment of FDR Control

Despite advances in DIA methodology, especially nDIA, the effectiveness of FDR control in such analyses remains insufficiently characterized(*26*). To address that, we used entrapment analysis to systematically evaluate the performance of FDR control across the search engines using entrapment analysis(*26*). Combined false discovery proportion (FDP) (upper bound) values were calculated for identifications filtered at a nominal 1% FDR.

First, we examined DDA results. In the mixed-species sample E5H50Y45, the FDPs of Tesorai Search remained below or close to 1% at both peptide and protein levels (Supplementary Figure 1A, B). FragPipe-DDA+ yielded higher FDPs compared to FragPipe DDA. The FDPs of FragPipe were significantly higher than 1% at peptide level but remained lower than 1% at protein level. This trend was similarly reflected in the HeLa DDA dataset (Supplementary Figure 1C, D), highlighting the relatively stronger FDR control offered by Tesorai Search in DDA workflows.

Turning to nDIA, Both DIA-NN and Tesorai Search maintained FDPs below 1% across both E5H50Y45 and HeLa datasets (Supplementary Figures 2A-D). The FDPs of FragPipe were higher than 1% at peptide level but remained lower than 1% at protein level. Overall, the FDPs observed for nDIA were on par with those from DDA with all three search engines, especially DIA-NN and Tesorai Search, reinforcing the reliability of nDIA identifications when stringent FDR control is applied.

### Quantitative Performance across Data Acquisition and Analysis Workflows

To assess the quantitative accuracy of different data acquisition and analysis workflows, we performed pairwise comparisons of MS signal intensities using sample pairs with known ratios of *E. coli* and yeast proteins in the mixed-species dataset. Specifically, we compared the following sample sets: E5H50Y45 vs. E45H50Y5 (ratio 9:1), E10H50Y40 vs. E40H50Y10 (ratio 4:1), and E20H50Y30 vs. E30H50Y20 (ratio 1.5:1), while human protein content remained constant across all samples (Figure 5A and Supplementary Figure 3A). Overall, DIA-based workflows demonstrated superior quantitative accuracy compared to DDA (Figure 5A-D and Supplementary Figure 3A-D). As expected, greater variation was observed at lower MS intensity levels (Figure 5A, and Supplementary Figure 3A).

**Figure 5.**
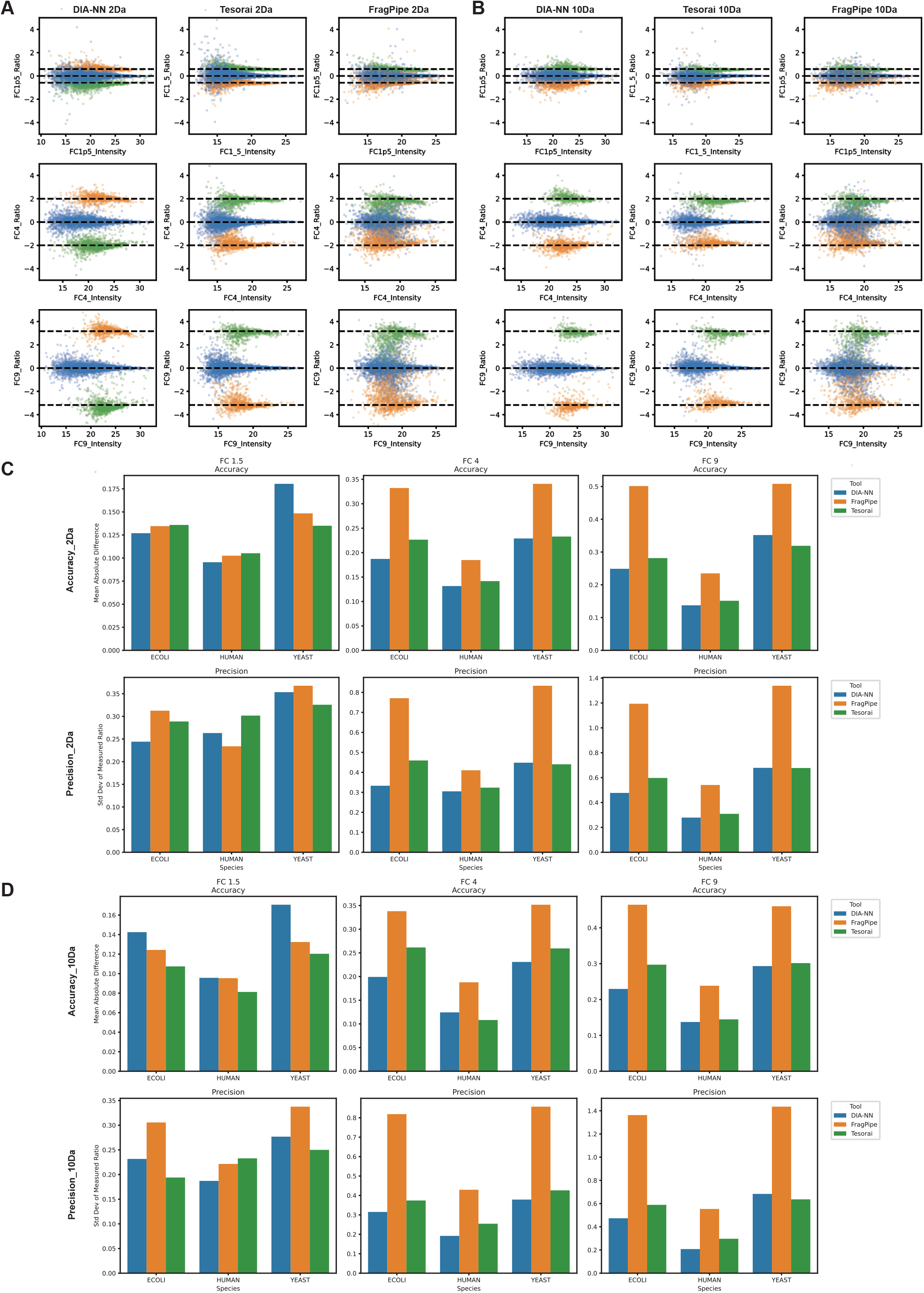
Quantitative performance in DIA data analysis. **(A, B)** Intensity and fold change of proteins identified from the benchmark samples by different search engines in DIA datasets with narrow **(A)** and wide **(B)** isolation window. Dashed lines represent expected log2(A/B) values for proteins from humans (green), yeast (purple), and *E. coli* (orange). FC_9, FC_4, and FC_1.5 are calculated from log2(E45H50Y5/E5H50Y45), log2(E40H50Y10/E10H50Y40), and log2(E30H50Y20/E20H50Y30), respectively. **(B)** Comparison of quantitative accuracy and precision for the output result of DIA-NN, Tesorai Search, and FragPipe from DIA datasets with narrow **(C)** or wide isolation windows **(D)**, respectively.

Among the DIA analysis tools, DIA-NN and Tesorai Search exhibited excellent quantitative performance (Figure 5A-D). DIA-NN, in particular, achieved the broadest dynamic ranges (Figure 5A). FragPipe exhibited limited quantitative accuracy with DIA data, despite its use of integrated DIA-NN modules for quantitation (Figure 5A-D). With DDA data, FragPipe standard DDA mode delivered more reliable quantitation than Tesorai Search (Supplementary Figure 3A-D), whereas FragPipe-DDA+ slightly impaired its quantitative performance.

In summary, among the tools evaluated, DIA-NN and Tesorai Search are well-suited for DIA-based quantitation, whereas FragPipe’s standard DDA workflow offers the most accurate quantitation for DDA datasets.

### High-Throughput nDIA-MS Confirms Target Depletion in Knockout Cell Lines

A common concern in biological research is that mass spectrometry (MS)-based proteomics often lacks quantitative rigor. In some cases, even substantial modulation of protein expression is not captured when analyte abundance is outside of the dynamic range. To assess the reliability of nDIA-MS for biological applications, we performed high-throughput, quantitative screening of a panel of 17 cancer-related KO cell lines, each targeting one protein involved in DNA damage response (DDR). All knockouts were previously validated by western blotting (Figure 6A and Supplementary Figure 4A).

**Figure 6.**
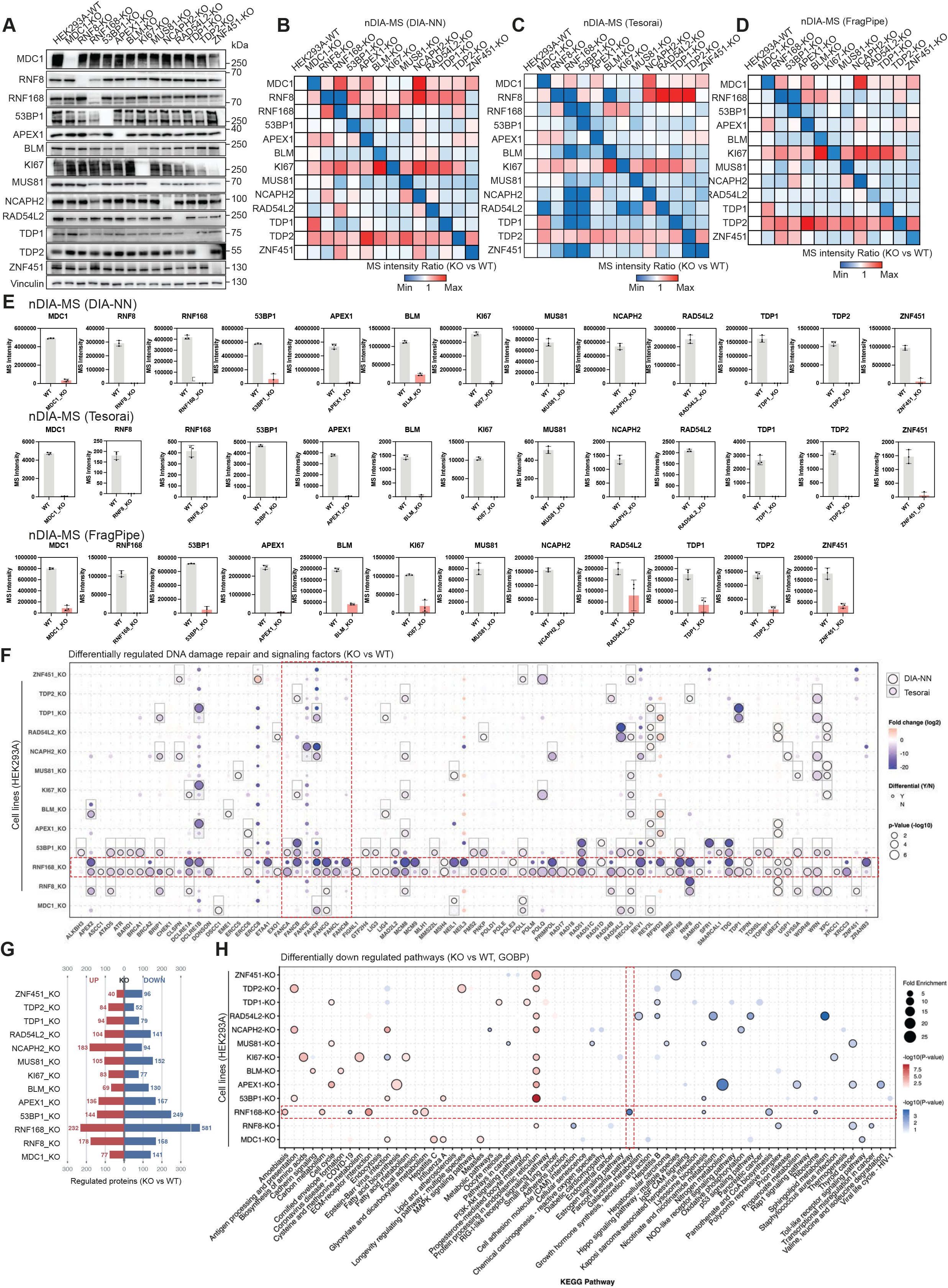
Label-free proteome profiling of knockout cell lines using nDIA-MS. **(A)** Western blotting results for the depletion of targeted proteins in knockout cell lines. Only the results of HEK293A cell lines were displayed. **(B-D)** Heatmap overview of the MS result for the abundance change of targeted proteins in knockout cell lines. Results were generated via **(B)** DIA-NN, **(C)** Tesorai Search, and **(D)** FragPipe, respectively. **(E)** nDIA-MS results confirm the downregulation of targeted proteins in knockout cell lines using all three search engines. **(F)** Profile of the DNA damage repair and signaling proteins that were differentially regulated in at least one of the tested knockout cell lines. **(G)** The number of differentially regulated proteins in each knockout cell line compared with wild-type cells. Only proteins identified by both DIA-NN and Tesorai Search were included. **(H)** Functional clustering of the differentially regulated proteins in the test HEK293A knockout cell lines. The result was plotted with KEGG pathway categories based on the DAVID database. Only top five categories with lowest p-value were included. Categories with p-values < 0.05 are indicated by black borders.

Based on the benchmarking results, nDIA was selected as the acquisition method. DIA data were subsequently processed using all three search engines (Figure 6B-D and Supplementary Figure 4B-D). DIA-NN successfully detected all 17 target proteins in WT samples (Figure 6B and Supplementary Figure 4B). Substantial downregulation of all target proteins was confirmed by MS, including 7 KO cell lines with complete loss of the target (Figure 6E and Supplementary Figure 4E). Tesorai Search also identified 16 of the 17 target proteins in WT samples, missing only RNF8 in HeLa cells (Figure 6C and Supplementary Figure 4C); in nearly all cases, target expression was not detected in the respective KO lines (Figure 6E and Supplementary Figure 4E). FragPipe identified 15 of the 17 target proteins in WT samples, with several proteins having less pronounced differences in expressions between WT and KO cell lines than the other two search engines (Figure 6D, E, and Supplementary Figure 4D, E).

After confirming target depletion, we assessed quantitative data for 10,224, 12,589, and 8,423 proteins across the tested cell lines using DIA-NN, Tesorai Search, and FragPipe, respectively (Supplementary Table 13), enabling system-wide insights generation across a broad range of functional protein networks and potential compensatory mechanisms (Figure 6B-D and Supplementary Figure 4B-D). For instance, MS analyses confirmed the interaction of proteins involved in the double-strand break (DSB) repair pathway, including MDC1, RNF8, RNF168, and 53BP1. Both DIA-NN and Tesorai Search revealed MDC1 was increased upon the knockout of RNF8 and RNF168 in HEK293A cells (Figure 6B, C). RNF8 was up-regulated in MDC1-KO and 53BP1-KO cells and down-regulated in RNF168-KO cells. RNF168 was up-regulated in RNF8-KO cells. Together, these results demonstrated that high-throughput nDIA-MS delivers the quantitative rigor needed for proteome-scale biological discovery.

### Proteomic Landscape of DDR Knockout Cell Lines

To further explore global DDR-related changes, we analyzed the DIA-NN and Tesorai Search output from all 17 of the HEK293A KO cell lines targeting individual DDR genes. DDR-associated proteins were curated based on a comprehensive genomic screen reported previously(*27, 28*). Across the dataset, 81 proteins showed significant modulation in at least one KO line (Figure 6F). Strikingly, the RNF168 KO line exhibited widespread downregulation of DDR proteins including several Fanconi anemia (FA) pathway proteins—FANCA, FANCB, FANCF, FANCG, FANCL, and FANCM (Figure 6F)(*29, 30*).

To better understand the biological implications of the observed effects, we performed functional clustering of all significantly regulated proteins in each KO cell lines (Figure 6G, H, Supplementary Table 14)(*31–34*). As anticipated, the Fanconi anemia pathway was specifically and significantly suppressed in RNF168 KO cells (p-value = 0.000409, fold enrichment = 5.79)(*33*). Additionally, the “protein processing in endoplasmic reticulum” category was enriched among the upregulated proteins in most KO cells. Closer examination of the proteins contributing to this enrichment revealed consistent upregulation of ER proteostasis factors across multiple KO cell lines, including HSPA5 (9/13 cell lines), HYOU1 (8/13), DNAJC3 (7/13), HERPUD1 (7/13), and PDIA4 (7/13), together with the stress-responsive chaperones CRYAB (6/13) and HSPH1 (4/13). These proteins are involved in ER protein folding, chaperone-mediated proteostasis, ER-associated degradation, and cellular stress responses, suggesting that loss of DDR factors may broadly perturb proteostasis and trigger an adaptive ER stress response(*35–38*). Together, these findings extend the functional footprint of DDR genes beyond DNA repair beyond canonical DNA repair pathways and demonstrate the utility of nDIA-MS for revealing previously unappreciated functional consequences of DDR perturbation.

## Discussion

The development of high-resolution, high-speed MS instruments enables narrow window DIA (nDIA) workflows to achieve unprecedented levels of throughput, enabling near-complete human proteome coverage in single-shot experiments(*8–10, 13*). nDIA now redefines the MS data acquisition and analysis toolkit^3^. Although DIA has been in development for decades, its widespread use was previously hindered by limited computational capacity to process the complex spectra it generates. That limitation led to the development of DDA, which selects precursor ions based on peak intensity and provides clear MS2 spectra for confident peptide and protein identification, but it omits many precursor (MS1) ions, decreasing proteome coverage and potentially introducing quantitative inaccuracies. In parallel, windowed DIA approaches, such as SWATH, use spectral libraries and fractionate precursor ions for fragmentation through stepped isolation windows (10–100 Th)(*39*). This decreases spectral complexity and omission of precursor ions, but the reliance on pre-generated spectral libraries from fractionated DDA runs remains a bottleneck, and the complexity of DIA spectra can still pose challenges for confident identification.

Recent advances in computational search engines, such as MSFragger-DDA+ and Chimerys, have improved the ability to deconvolute chimeric spectra and confidently identify co-fragmented peptides(*18, 40*). Building on those developments, wide-window acquisition (WWA) was recently introduced as a modified DDA approach with isolation windows ranging from 5 to 20 Th(*14*). Although WWA enhances DDA protein identification sensitivity, it still suffers from the specificity (at least at the peptide level), as well as quantitative limitations of DDA where only high-intensity peaks are selected for fragmentation. The Orbitrap Astral system, with a high scan speed of 200 Hz and high mass resolution of ~80,000, enabled the development of nDIA(*6*). Narrow isolation window, such as 2 Th, decreases MS2 spectral complexity but requires rapid acquisition of hundreds of stepped windows across the mass range of peptides (typically from 100 to 2000 m/z for tryptic peptides), rendering the approach impractical for conventional MS instruments but manageable with ultra-high scan speed(*3*).

In this study, we compared the performance of nDIA, wide-window DIA (10-100 Th), conventional DDA (0.5-2 Th) and wide-window acquisition (WWA, 5-20 Th) strategies(*14*) using peptide mixtures from *E. coli*, human, and yeast in various ratios(*9*). We analyzed those datasets using Tesorai Search, a pre-trained model-based search engine capable of processing both DIA and DDA datasets and compared it to DIA-NN and FragPipe(*15–17*). Our results showed that nDIA outperformed other strategies in proteome coverage and quantitative accuracy. The combination of nDIA with DIA-NN and Tesorai Search provides the greatest depth of proteome coverage. FragPipe excelled at processing DDA datasets. Specifically, the FragPipe-DDA+ workflow significantly improved protein identification rates compared to the conventional DDA workflow. Moreover, the DDA+ workflow produced substantial overlap of protein IDs with DIA, providing evidence of mutual validation between the two approaches(*18*).

The emphasis on protein identification depth without adequate validation of identification accuracy is a persistent challenge in the field(*26*). Although standard target-decoy competition is theoretically sufficient for FDR control, most modern search engines for peptide annotation employ on-the-fly training of AI/ML classifiers to distinguish targets from decoy peptide sequences. This practice introduces risks of overfitting, as the same target-decoy dataset is effectively used twice—both for training and evaluation. Additionally, features used during training may inadvertently encode information that differentiates targets from decoys, leading to biased FDR estimates. Among the tools benchmarked in this study, Tesorai Search(*15*) is the only engine that relies on a pre-trained classifier and avoids on-the-fly model training. It is encouraging that this approach performs well in both identification depth and accuracy—especially considering that the model in Tesorai Search was not specifically optimized for Orbitrap Astral MS data. These findings validate the robustness of the pre-trained approach and pave the way for how search engines could be designed for even greater rigor and reproducibility in the future.

To demonstrate its performance, we applied nDIA-MS to screen 17 cancer cell lines with single knockouts of specific DNA damage response genes(*27*). The MS data confirmed significant downregulation of the targeted proteins and revealed functional consequences of DDR gene loss beyond DNA repair, including extensive suppression of proteins involved in Fanconi anemia (FA) proteins following RNF168 depletion(*20, 41, 42*) and recurrent upregulation of ER proteostasis factors across multiple KO cell lines(*37, 38*). These results highlight the value of nDIA-MS proteomics for large-scale biological investigations and uncover previously unrecognized consequences of DDR gene perturbation.

## Materials and method

### Cells and cell culture

HeLa cells were purchased from the American Type Culture Collection (ATCC) (Manassas, VA) and maintained in Dulbecco’s modified Eagle’s medium containing 10% fetal calf serum at 37°C with 5% CO2. HEK293A cells were purchased from Thermo Fisher Scientific (R70507) and maintained in Dulbecco’s modified Eagle’s medium containing 10% fetal calf serum at 37°C with 5% CO2.

### Chemicals and reagents

Water (LC-MS grade, Optima, Cat. No. 10509404), acetonitrile (LC-MS grade, Optima, Cat. No. 10001334), and formic acid (LC-MS grade, Thermo Scientific Pierce, Cat. No. 13454279) were purchased from Fisher Chemicals. MassPREP E. coli digest standard (Cat. No. 186003196) was purchased from Waters. HeLa protein digest standard (Thermo Scientific Pierce, Cat. No. 88328) and yeast digest standard (Thermo Scientific Pierce, Cat. No. A47951) were purchased from Thermo Scientific. Sequence grade trypsin (Cat. No. V5113) were purchased from Promega.

### Western blotting

Cells were washed with PBS and resuspended in 1×Laemmli buffer for cell lysis. The lysate was then boiled at 95°C for 10 minutes and analyzed by Western blotting.

### MS sample preparation

5 × 10^6^ cells from each sample were washed twice with 1x PBS, then centrifuged at 1,000 rpm for 3 min at 4°C. The collected pellets were resuspended in ice-cold extraction buffer containing 4 M urea and 50 mM NH_4_HCO_3_. The samples were then sonicated using a microtip sonicator at 35 W for 2 × 10 s pulses. Homogenates were clarified by centrifugation at 10,000 × g for 10 min at 4°C. Protein concentrations were determined using the BCA assay (Thermo, Pierce). Proteins in the supernatant were denatured by boiling at 95°C for 5 min. For mass spectrometry (MS) sample preparation, 100 µg of each protein sample was diluted to a final volume of 100 µL with 50 mM NH_4_HCO_3_. The samples were then reduced with 5 mM DTT at 37 °C for 1 h, followed by alkylation with 15 mM iodoacetamide at room temperature in the dark for 30 min. The reaction was quenched with an additional 15 mM DTT. Proteolytic digestion was performed by adding 5 µL of 400 ng/µL trypsin (Promega) and incubating the samples at 37°C overnight. The samples were acidified with 1 µL of 10% formic acid (final concentration ~0.1% v/v) and centrifuged at 10,000 × g for 10 min at 4°C. The supernatant was desalted using a BioPureSPN Mini PROTO 300 C18 column (The Nest Group, Cat. No. HUM S18V), dried in a vacuum, and stored at −80°C until further analysis.

### LC-MS/MS analysis

LC-MS/MS analyses were performed using Orbitrap Astral mass spectrometer coupled with Vanquish Neo UHPLC system (Thermo Fisher Scientific). The vacuum dried peptide samples were resuspended with 0.1% of formic acid. Peptides were separated using a C18 column (CoAnn Technologies, Cat. No. HEB07502001718I, 75 μm × 20 cm) at a flow rate of 400 nL/min. Peptides were chromatographically separated using a linear gradient of solvent B (0.1% formic acid in ACN) and solvent A (0.1% formic acid in water) unless otherwise specified. Linear gradients were as follows: from 2% to 8% of solvent B 1.5 min, 8% to 38% of solvent B from 1.6 to 12.6 min, 38% to 100% of solvent B from 12.6 to 13 min, 100% B from 13 to 15 min. For DIA analysis on the Orbitrap Astral MS, MS1 spectra were collected in the Orbitrap every 0.6 s at a resolution of 240,000. The full scan range was 380-980 m/z. The MS1 normalized AGC target was set to 500% with a maximum injection time of 50 ms. DIA MS2 scans were acquired in the Astral analyzer over a range of 380-980 m/z with a normalized AGC target of 500% and a maximum injection time of 2.5 ms and an HCD collision energy setting of 25%. The isolation window was set at 2 Th or 10 Th without window overlap unless otherwise specified. For DDA analysis, the data dependent mode was set at cycle time. The time between master scans was set at 0.6 s. Other parameters were similar to DIA analysis.

### MS data analysis

Raw files from DIA experiments were analyzed using DIA-NN 2.0 Academia, Tesorai Search, and FragPipe. For DIA-NN analysis, the in-silico spectral library was generated from a combined three-species databased including sequences from the human reference database (UP000005640, UniProt 2024 release, 20,654 entries), the Escherichia coli reference database (UP000000625, UniProt 2024 release, 4,402 entries), the yeast reference database (UP000002311, UniProt 2024 release, 6,065 entries) allowing N-term M excision and 1 missed cleavage. The DIA-NN search included the following settings: Protein inference = ‘Genes’, Neural network classifier = ‘Single-pass mode’, Quantification strategy = ‘Robust LC (high precision)’, Cross-run normalization = ‘RT-dependent’, Library Generation = ‘Smart Profiling’ and Speed and RAM usage = ‘Optimal results’. Mass accuracy and MS1 accuracy were set to 0 for automatic inference. ‘No share spectra’, ‘Heuristic protein inference’ and ‘MBR’ were checked. The output results from DIA-NN were filtered for Q.Value <=0.01 and PG.Qvalue <=0.01. For Tesorai Search analysis, the parameters were set as follows: allowed missed cleavages = ‘2’, digestion enzyme = ‘Trypsin_P’, fixed modification = ‘Carbamidomethyl_C’, variable modification = ‘Oxidation_M, Acetylation_Nterm’, max peptide length = ‘50’, min peptide length = ‘7’. For FragPipe analysis. The search type is set at DIA or DDA depending on the data acquisition approach of the raw files. For FragPipe-DIA analysis, the default settings of the ‘DIA-SpecLib-Quant’ workflow was employed. The raw files were converted to mzML files using MSConvert. MSFragger was used for identification. MSBooster was employed. Spectral library generation from search results was enabled. The integrated DIA-NN 1.8.2_beta_8 module was employed for quantification. For FragPipe-DDA analysis, the default setting of the ‘LFQ-MBR’ workflow was employed. IonQuant 1.10.27 was used for quantification. ‘Match between runs’ was checked. For FragPipe-DDA+ analysis, the default setting of the ‘WWA’ workflow was employed. The data type was set to DDA+.

### Tesorai DIA search

The DIA extension of the Tesorai Search workflow builds on the same architecture and principles as its DDA counterpart(*43*), but is adapted to operate on extracted ion chromatograms (XICs) rather than individual MS2 scans. For each candidate peptide-spectrum match, theoretical fragment m/z values are computed and used to extract XICs within a fixed ±10 retention time window around the chromatographic apex. These XICs form an image-like spectrum representation, while the modified peptide sequence is embedded into a vector form via the same sequence encoder used in the DDA model. The peptide and XIC representations are then concatenated and passed through a joint encoder that outputs a single score reflecting match quality. This model was pre-trained once using approximately 100 million DIA XICs–PSM pairs drawn from Astral and diverse standard DIA datasets spanning multiple species. No decoy sequences or dataset-specific retraining are required, enabling robust peptide identification directly from DIA data using the same scalable and inference-efficient infrastructure as in the Tesorai-DDA workflow.

### Entrapment analysis

To assess FDR control in proteomics data analysis, we used an entrapment strategy by adding proteins to the search database that are known to be absent from the sample. These entrapment proteins act as internal negative controls for estimating false discovery proportions (FDP). For the 3-mix experiments, we combined FASTA files from Human, E. coli, and Yeast (31,121 proteins total) with Ricinus communis (Castor bean; 31,219 proteins) as the entrapment species. For the HeLa experiments, the Human FASTA (20,654 proteins) was similarly combined with R. communis. We also included common contaminant proteins. To avoid inflating false discovery estimates, any entrapment or contaminant protein that shared at least one peptide with a target protein was excluded from the entrapment set. Sample-level FDP (lower bound) was calculated as the number of identified entrapment proteins divided by the total number of proteins identified in that sample (including both target and entrapment proteins). Combined FDP (upper bound) was calculated by adjusting for the relative size of the entrapment and target databases, as previously described(*26*).

## Acknowledgments

We thank all members of MD Anderson Proteomics Core Facility and Metabolomics Core Facility for their help and constructive discussions.

## Author contributions

Conceptualization: Y.X. and P.L.L. Methodology: Y.X., D.S., and H.Z. Investigation: Y.X., H.Z, L.T, and B.W. Data Processing: Y.X. and D.S. Visualization: Y.X. Supervision: P.L.L, P.C. and J.N.W. Writing—original draft: Y.X. Writing—review and editing: Y.X., H.Z., D.S. P.L.L, P.C. and J.N.W.

## Data and materials availability

The mass spectrometry proteomics data generated in this study have been deposited to the ProteomeXchange Consortium via the MassIVE repository with the dataset identifier PXD066228 (MassIVE ID: MSV000098543).

## Supplementary Materials

Supplementary Figures 1-4

Supplementary Tables 1-14

